# Chemophototherapy Overcomes Doxorubicin Resistance in Human Ovarian Cancer Cells

**DOI:** 10.1101/2022.02.14.480412

**Authors:** Sanjana Ghosh, Upendra Chitgupi, Ulas Sunar, Jonathan F. Lovell

**Author notes:** Corresponding author: Jonathan F. Lovell, University at Buffalo, State University of New York, 210 Bonner Hall, Buffalo 14260.

## Abstract

Porphyrin-phospholipid (PoP) liposomes loaded with Doxorubicin have been demonstrated to be an efficient vehicle for chemophototherapy (CPT). Multi-drug resistance (MDR) of cancer cells is a known resistance mechanism for cancer chemotherapies. We report a phototherapeutic measure to overcome Dox-resistance using Doxorubicin (Dox)-loaded PoP liposomes. In vitro studies using free Dox or Dox loaded into liposomes with 2 mol.% showed human ovarian carcinoma A2780 cells were more susceptible to these drugs compared to the corresponding Dox-resistant A2780-R cells. In contrast, when CPT was applied with LC-Dox-PoP liposomes, effective killing of both non-resistant and resistant A2780 cell lines was observed. An *in vivo* study to assess the efficiency of LC-Dox-PoP liposomes with phototreatment showed effective tumor shrinkage and prolonged survival of athymic nude mice bearing A2780-R tumor xenografts. Biodistribution analysis demonstrated enhanced tumoral drug uptake in Dox-resistant tumors with CPT, pointing to the likelihood that increased drug delivery overcame resistance mechanisms to provide for improved anti-cancer therapy.

## 1. Introduction

Ovarian cancer is one of the most lethal type of gynecological cancers and frequently develops without showing detectable symptoms so is diagnosed at an advanced stage [1]. The prognosis for ovarian cancer patients is poor with a 5-year survival rate of 3-19% for patients diagnosed at stage III or IV. If patients are diagnosed at an early stage, they have a 5-year survival rate of 40-90%, consequently, efforts are being focused on improving therapeutic techniques to cure patients detected by early diagnosis [2]. The standard of care for advanced ovarian cancer involves surgery to reduce tumoral mass followed by combination chemotherapy with platinum and taxane drugs [3, 4]. Doxorubicin (Dox) is an anthracycline antibiotic with a broad spectrum of chemotherapeutic competence [5, 6]. Although chemotherapy is a well-established modality, a high percentage of patients previously treated with chemotherapy show tumor recurrence and need further therapies [7, 8]. Pegylated liposomal Doxorubicin (Doxil in the USA or Caelyx in Canada and Europe) was the first liposomal chemotherapy drug to be FDA approved in 1999 and by the European Medicines Evaluation Agency (EMA) in 2000 as a monotherapy for the treatment of advanced ovarian cancer patients following failure of first-line treatment with platinum-taxane combination.

Many human cancers, including ovarian cancer, have been found to develop resistance to multiple chemotherapeutic drugs over time. This resistance to cytotoxic drugs, also referred to as pleiotropic resistance or multi-drug resistance (MDR), limits chemotherapy efficacy for cancer treatment. MDR is usually associated with reduced tumoral drug accumulation and increased removal of the drug from the tumor. Reappearance of tumor cells after eradication of cancer cells by chemotherapy is usually attributed to the growth of cancer cells with developed resistance to a previously administered drug or a group of drugs. The effect of any chemotherapeutic drug on MDR cells has been observed to be reduced even if the drug possesses a chemically functional group or mechanism of action different from the previously administered drugs [9–22]. Furthermore, MDR has been associated with overexpression of ATP-binding cassette (ABC) transporters such as P-glycoproteins (P-gp), multidrug-resistance protein (MRP), and breast cancer resistance protein (BCRP) on the surface of cancer cells [23–26]. Specifically, P-gp overexpression has been reported to be one of the causes of MDR in ovarian cancer [27, 28]. P-gp is a transmembrane protein that mediates cellular efflux of its substrates and many chemotherapeutic agents including Dox [29].

To overcome MDR, numerous approaches have been studied. Targeted drug delivery to cancer cells that can prevent tumoral drug efflux could be an effective approach for this. This may be achieved by encapsulation or attachment of chemotherapeutic drugs to nanoparticles made of polymers or lipids [30–35]. Drug delivery by encapsulation of drugs inside liposomes is thought to be a potential strategy to curb MDR [36]. Improved drug delivery to cancer cells beyond a certain threshold could also overcome MDR.

Liposomes are bilayered nanostructures made up of phospholipids and are of utility for photosensitizer delivery [37, 38]. Liposomal PDT formulations have been investigated for ovarian cancer research [39]. Targeted drug delivery using liposome triggered-release techniques has been reported to be effective in augmented tumoral drug uptake resulting in enhanced therapeutic results [40, 41]. These techniques include strategies that depend on heat [42, 43], pH [44, 45], enzymes [46–48], magnetic pulses [48], and light [49, 50]. Near-infrared (NIR) light is known to be harmless and can penetrate tissue beneath the surface. This makes NIR light fascinating as a liposomal trigger technique for faster and localized drug release [51–55]. As a result, numerous studies have recently been conducted to develop light-triggered nano-vehicles [56–58]. Dox entrapped in liposomes has been found to have enhanced therapeutic effects in tumor xenograft models over free drug. These effects include improved tumoral drug uptake by enhanced permeability and retention (EPR) effect and increased circulation time in the blood, as well as decreased cardiac toxicity [59–61]. In addition, it has been reported to effectively annul the MDR phenotype [40, 62, 63].

Recently, a photosensitive liposomal formulation of Dox with prolonged blood circulation time was reported. This formulation was developed by adding 2 mol% porphyrin-phospholipid (PoP) to the FDA-approved liposomal formulation of Dox, DOXIL. PoP is a photosensitive agent that can be used to create stable photoactivatable liposomes that release the drug upon light irradiation [64]. In this study, we investigate the impact of LC-Dox-PoP liposomes on the overcoming of Dox resistance and the therapy of cancer.

## 2. Materials and Methods

### 2.1. Materials

All lipids were acquired from Corden Pharma International, and other materials were acquired from Sigma, if not mentioned otherwise. Porphyrin phospholipid or PoP was synthesized as described before [65]. As reported earlier, the formulation for LC-Dox-PoP liposomes included 53 mol. % 1,2-distearoyl-sn-glycero-3-phosphocholine (DSPC, Corden Pharma# LP-R4-076), 40 mol. % cholesterol (PhytoChol, Wilshire Technologies Inc. #57-88-5), 2 mol. % PoP and 5 mol. % 1,2-distearoyl-sn-glycero-3-phosphoethanolamine-N-[methoxy(polyethylene glycol)-2000] (MPEG-2000-DSPE, Corden Pharma# LP-R4-039) [64].

### 2.2. Liposome preparation and drug loading

To prepare 5 mL (20 mg/mL) of LC-Dox-PoP liposomes, 100 mg of lipids as per the mentioned ratio was dissolved in 1 mL ethanol at 60 °C and then 4 mL of 250 mM ammonium sulfate (pH 5.5) was injected into the lipid mixture. The lipid mixture was then extruded 10 times at 60-70 °C through sequentially stacked polycarbonate membranes of 0.2, 0.1 and 0.08 μm pore size in a high pressure nitrogen extruder (Northern lipids), followed by dialysis (at least twice) in 800 mL solution of 10% sucrose with 10 mM Histidine (pH 6.5) buffer to remove free ammonium sulfate. Dox (Dox LC Labs # D-4000) was loaded in the liposomes by incubating Dox in PoP liposomes at 60 °C for 1 hour in Dox:lipid loading molar ratio of 1:5.

### 2.3. Cell culture and maintenance of non-resistant and Dox resistant A2780 cells

Human ovarian cancer A2780 and Adriamycin resistant human ovarian cancer A2780-R cells were procured from Sigma Inc. Cells were subcultured in RPMI 1640 media with no folic acid supplemented with 10% fetal bovine serum and 1% antibiotics. For in vitro studies, cells were seeded in a 96-well plate at 2×10^5^ cells per well. The 96-well plate was placed in an incubator maintained at 5% CO_2_ and 37° C.

### 2.4. Dark-toxicity in vitro studies

Cells seeded in a 96-well plate were incubated with either free Dox or Dox-loaded PoP liposomes. Cells were incubated with samples at the indicated concentrations for 24 hours and were then washed twice with PBS to remove free dox and liposome samples in the media. Cells were incubated in RPMI 1640 containing serum for 24 hours, followed by the XTT cell viability assay to assess dark toxicity.

### 2.5. Chemophototherapy in vitro studies

A2780 and A2780-R cells were seeded in a 96-well plate and treated with either dox alone or LC-Dox-PoP liposome sample. Cells were washed with PBS twice followed by the addition of fresh RPMI media supplemented with FBS. 96-well plate was placed under a laser box equipped with 665 nm laser light. A powermeter was used to measure the power generated by the box. Power irradiation was set to 23.5 mW/cm^2^ for 7 minutes to impart 10 J/cm^2^ energy and 14 minutes for 20 J/cm^2^. The plate was not exposed to light, which served as control and is indicated by 0 J/cm^2^ in **Fig. 3**.

### 2.6. Tumor growth inhibition

All murine tumor model experiments were conducted in accordance with the standard protocols of University at Buffalo’s Institute of Animal Care and Use Committee (IACUC). Five-week-old female athymic nude mice were injected with 2 × 10^6^ A2780-R cells (Dox resistant cells) subcutaneously on the right flank near the groin. When the tumors grew about 4-6 mm in diameter, mice bearing A2780-R tumors were randomly grouped into 3 cohorts. Mice that received treatment were intravenously injected with 4 mg/kg LC-Dox-PoP liposomes. Mice that received laser treatment were treated with a 665 nm laser diode (RPMC laser, LDX-3115-665) at a fluence rate of 200 mW/cm^2^ at a total laser fluence of 250 J/cm^2^. Tumor size and health of mice were monitored 2-3 times per week and tumor volumes were calculated using the ellipsoid formula: Volume = π.L.W.H/6, where L, Wand H are the length, width, and height of the tumor, respectively. The mice were sacrificed when the tumors grew to about 1.5 cm in diameter.

### 2.7. Biodistribution

Female athymic nude mice (~5 weeks old) were inoculated with 2 × 10^6^ A2780 cells on both flanks. When the tumor size reached 6-8 mm (n = 4) they were administered with 4 mg/kg LC-Dox-PoP liposomes via tail-vein intravenous injection. Post 1 hour from drug administration, one tumor from each mouse was treated with 200 mW/cm^2^ with a total fluence of 200 J/cm^2^ using a 665 nm laser diode (RPMC laser, LDX-3115-665). All mice were sacrificed post 8 hours from drug administration. Tumor and key organs were harvested. To determine the accumulation of the drug in tissues, ~100 mg of tissue was homogenized in nuclear lysis buffer [0.25 mol/L sucrose, 5 mmol/L Tris HCl, 1 mmol/L MgSO_4_,1 mmol/L CaCl_2_ (pH 7.6)] and extracted overnight in 0.075N HCI 90% isopropanol. The amount of Dox and PoP was determined by fluorescence measurements.

## 3. Results and Discussion

### 3.1. Characterization of LC-Dox-PoP liposomes

The active loading of Dox into PoP liposomes via ammonium sulfate gradient was found to be more than 98% of the Dox added based on gel filtration analysis (**Fig. 1A**). The size and PDI of the liposomes post loading were found to be 112.3 ± 1.3 nm and 0.04, ± SD 0.013, respectively. To find the drug release kinetics, the liposomes were laser-irradiated while stirring in a custom set-up in a fluorometer under a 665 nm diode laser at a fluence of 300 J/cm^2^ at 37 °C in 50% bovine serum. More than 95% of the encapsulated drug was released within 5 min of laser irradiation. The serum stability test of the LC-Dox-PoP liposomes in 50% bovine serum at 37 ° C showed almost negligible release over 6 hours. The spectra of empty PoP liposomes (**Fig. 1B**) and LC-Dox-PoP liposomes (**Fig. 1C**) were recorded on a fluorometer diluted in ethanol at 200x dilution and the drug concentrations were calculated using standard curves. Following Dox loading, the PoP concentration was 0.5 mg/mL and Dox concentration was 2.9 mg/mL.

**Figure 1:**
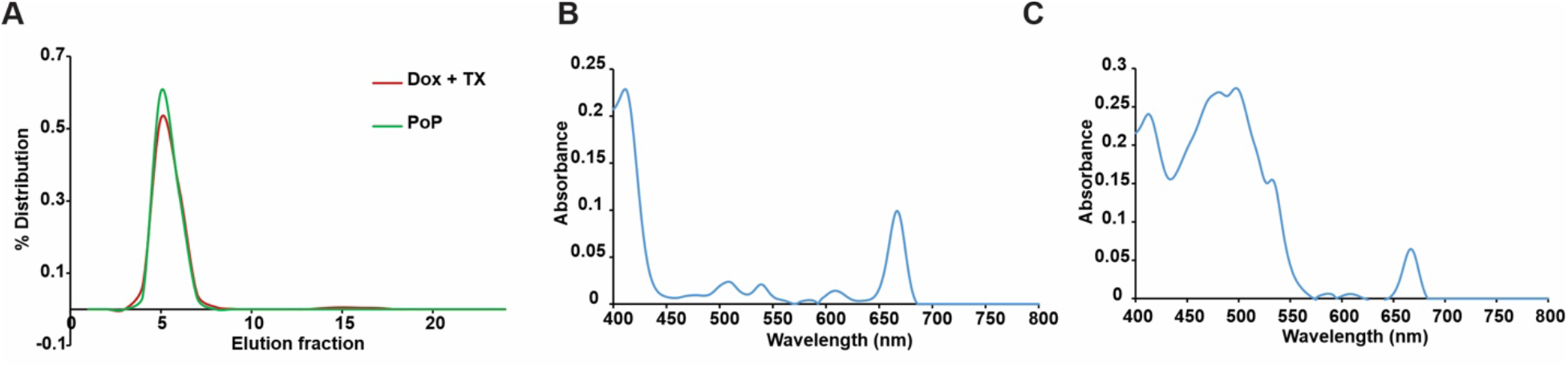
Characterization of LC-Dox-PoP liposomes. 100 mg of PoP liposomes were prepared and Dox was loaded into the liposomes at a loading ratio of 1:5 drug:lipids. Post loading, characterization of liposomes was done as mentioned in the methods section to maintain quality control and batch record was documented. **A)** Loading of Dox (Dox + TX shows the loading of Dox inside the liposomes when treated with 10% Triton X detergent) **B)** Spectra of empty PoP liposomes **C)** Spectra of LC-Dox-PoP liposomes.

### 3.2. Effect of Dox and LC-Dox-PoP liposomes on A2780 and A2780-R cells

Human ovarian cancer cells (A2780) and Dox-(also termed Adriamycin) resistant human ovarian cancer cells (A2780-R) seeded in a 96-well plate were incubated with either free Dox or LC-Dox-PoP liposomes containing different Dox concentrations (0.5 μg/mL, 1 μg/mL, 5 μg/mL, 10 μg/mL and 50 μg/mL Dox) for 24 hours and then washed with PBS twice to remove free dox and liposomes in the media. After incubating the cells in RPMI 1640 (with serum) for 24 hours, the XTT cell viability assay was carried out to assess dark toxicity. **Fig. 2** shows the results of the cell viability assay done by the XTT assay. **Fig. 2A** shows the effect of Dox alone on the cells. At the highest Dox concentration of 50 μg/mL, the percentage of viable A2780 cells was only approximately 20% of the total number of cells, whereas the percentage of viable A2780-R cells was greater than 60% of the total number of cells. **Fig. 2B** shows the effect of LC-Dox-PoP liposomes on the cells. At the highest Dox concentration of 50 μg/mL, the percentage of viable A2780 cells post 24 hours of incubation was about 30% of the total number of cells, whereas the percentage of viable A2780-R cells was above 74% of the total number of cells. **Fig. 2C** shows the effect of LC-Dox-PoP liposomes with irradiation on the cells. At a Dox concentration of 25 μg/mL, the percentage of viability for both types of cells, the non-resistant A2780 cells and the Dox-resistant A2780-R cells was only about 17% of the total number of cells. This shows the impact of irradiation on resistant A2780 cells. Therefore, this result adequately shows that laser irradiation at a total fluence of 20 J/cm^2^ overcomes the resistance of A2780-R cells to Dox, resulting in significantly increased cell death that was not observed in the experiments with Dox alone or LC-Dox-PoP liposomes containing the same Dox concentrations without irradiation.

**Figure 2:**
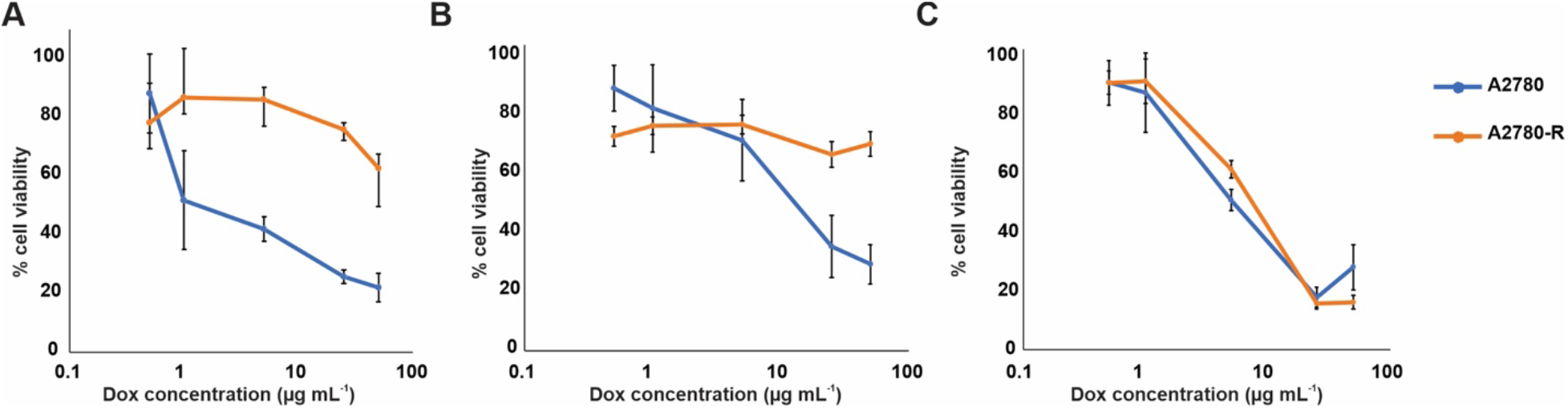
Dox-dependent cell death. A2780 ovarian cancer cell line and A2780-R dox resistant ovarian cancer cell line incubated with **A)** Dox alone **B)** LC-Dox-PoP liposomes **C)** LC-Dox-PoP liposomes followed by 665 nm laser irradiation at 20 J cm^-2^.

### 3.3. Impact of phototherapy on A2780 and A2780-R cells

To study the impact of CPT, human ovarian cancer cells (A2780) and Adriamycin resistant human ovarian cancer cells (A2780-R) cells seeded in a 96-well plate were treated with PoP-liposomes empty or loaded with Dox or free Dox and then laser treated using a laser box equipped with 665 nm laser light with laser fluence rate of 23.5 mW cm^-2^ for 7 minutes to impart 10 J cm^-2^ energy and 14 minutes for 20 J cm ^-2^. **Fig.3** demonstrates the effect of laser on Dox resistance of A2780-R cells. **Fig.3A** shows the effect of the laser when the cells were treated with 25 ug/mL Dox alone and irradiated with different laser powers. The percentage of viable cells from A2780 cells dropped to 20% without irradiation and was generally similar with irradiation. On the other hand, the percentage of viable cells of A2780-R cells was found to be about 80% without irradiation and stays above 65% with irradiation of 10 J/cm^2^ laser fluence. However, the percentage cell viability of A2780-R cells was found to be around 60% with a higher laser power of 20 J/cm^2^. The increase in cell death could be the result of photothermal injury to cells. **Fig. 3B** shows the effect of empty-PoP liposomes with irradiation. The percentage of viable cells of both cell types was found to be 100% with no irradiation. This shows that the PoP-liposomes did not have toxicity toward the cells. The percentage of viable cells of both cell types, at 10 J/cm^2^, drops to around 10%, which drops a little more at laser power 20 J/cm^2^. **Fig. 3C** shows the effect of LC-Dox-PoP liposomes with irradiation. It was observed that 25 ug/mL of Dox in LC-Dox-PoP liposomes with no irradiation was able to kill about 80% of the non-resistant A2780 cells, while the cell viability of A2780-R cells was found to be as high as 93%. At irradiation of 10 J/cm^2^, it was observed that the cell viability of both cell types dropped to about 10%. This shows the impact of photochemotherapy on overcoming the Dox-resistance of A2780-R cells.

**Figure 3:**
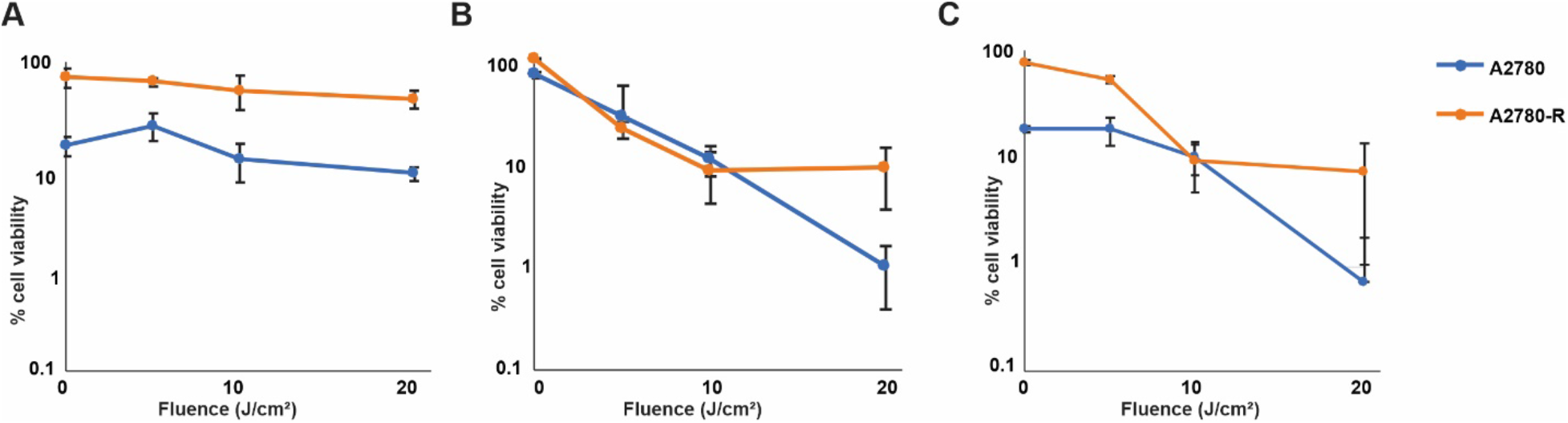
Phototherapy of Dox-resistant A2780 cells. A2780 and A2780-R cells were treated with (A) Dox (B) empty PoP and (C) Dox-PoP liposomes followed by irradiation with a 665 nm laser at varying fluence. Mean ±SD for n=3 datapoints obtained at each timepoint.

### 3.4. Phototherapy overcomes Dox-resistance in tumors

To test the efficacy of chomophototherapy in curbing Dox-resistance in vivo, a tumor inhibition study was conducted. Mice with A2780-R tumors were untreated or intravenously injected with LC-Dox-PoP liposomes containing 4 mg/kg Dox dose. After 1 hour from drug injection, mice that received laser treatment were treated with a 665 nm laser diode (RPMC laser, LDX-3115-665) at a fluence rate of 200 mW/cm^2^ at a total laser fluence of 250 J/cm^2^. The relative tumor volume growth data over time demonstrate the tumor-inhibiting capability of chemophototherapy using LC-Dox-PoP liposomes, as shown in **Fig. 4A**. It was observed that the tumors treated with LC-Dox-PoP liposomes with laser irradiation reduced steadily with time whereas tumors from other cohorts grew rapidly. By the 10^th^ day from treatment, the tumors of mice treated with photochemotherapy were reduced to less than half of their initial size, whereas the tumors of mice from other groups increased to >5X their initial size. Furthermore, the percentage of mice that survived without reaching the endpoint of tumoral diameter approved in the protocol provided by IACUC was plotted using a Kaplan Meier curve, as shown in **Fig. 4B**. It was observed that all the untreated mice had to be sacrificed by the 27^th^ day of the study because the tumors grew more than 1.5 cm in diameter (the endpoint approved by IACUC). The mice treated with LC-Dox-PoP liposomes with no irradiation reached their endpoint within 33 days from treatment. This shows that there was little effect of chemotherapy by LC-Dox-PoP liposomes accumulated in tumors that received no laser treatment, as shown by delayed average tumor volume growth of the cohort by about 6 days over untreated mice. On the other hand, the mice treated with CPT showed significantly faster tumor shrinkage and they survived throughout the time of the study without further tumor recurrence. The improved and faster shrinkage of tumors by CPT distinctly demonstrates the impact of phototherapy on Dox-resistant tumors. This enhanced therapeutic effect in tumor ablation by CPT is probably due to the increased accumulation of Dox in tumors as a result of better penetration of Dox through damaged tumor vasculature attributed to phototherapy [64].

**Figure 4:**
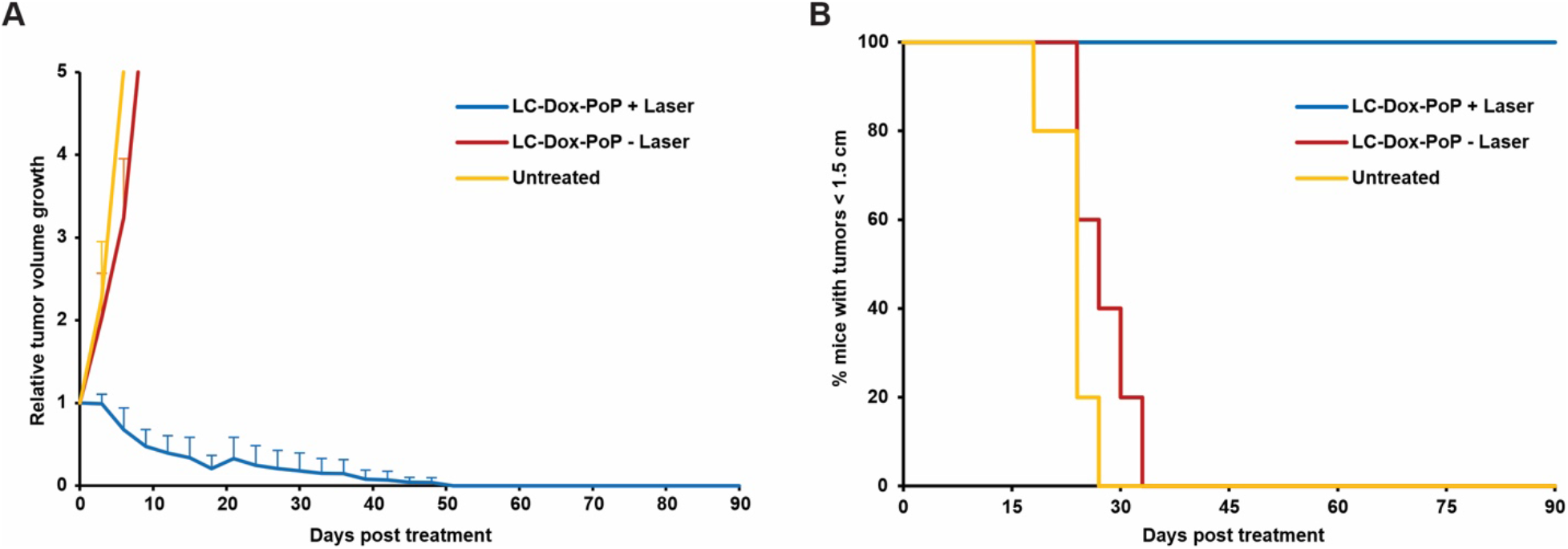
Tumor growth inhibition of Dox-resistant A2780 cells. Mice with A2780-R cells untreated/treated with LC-Dox-PoP liposomes with 4 mg/kg Dox. Mice with laser treatment were treated using a 665 nm laser with a laser fluence rate of 200 mW/cm^2^ with a total fluence of 250 J/cm^2^. (A) Relative tumor growth data (B) %Survival curve over time. Each data point shows the mean and ±SD of n=5 mice data obtained at each time point.

### 3.5. Tumoral drug accumulation with chemophototherapy

To assess the reason behind enhanced tumoral inhibition in Dox-resistant tumors by chemophototherapy, a biodistribution study of Dox and PoP in tumors and key organs was conducted. A dual tumor model of female athymic nude mice bearing A2780 human ovarian cancer xenografts was treated with LC-Dox-PoP liposomes with a 4 mg/kg Dox dose. After 1 hour from drug administration, one tumor of each mouse was treated with a 665 nm laser with a laser fluence rate of 200 mW/cm^2^ with a total fluence of 250 J/cm^2^. All mice were sacrificed post 8 hours from drug administration and tissues were harvested. The drug accumulation in tumors, liver, kidney, spleen, and heart were analyzed. **Fig. 5B** and **5C** show the biodistribution of Dox and PoP in liver, kidney, spleen and heart. The study demonstrates an almost negligible accumulation of Dox in tumors without irradiation, whereas tumors with irradiation show a mean of about 3 μg/gm Dox in them, as shown in **Fig. 5A**. It also shows the deposition of a small amount of PoP in tumors without irradiation, whereas tumors with irradiation show an average of about 1.5 ug/gm PoP in them. The distinctly enhanced deposition of Dox and PoP in tumors treated with laser over tumors untreated with irradiation clearly indicates that phototherapy can lead to significantly improved distribution of drugs in tumors and key organs. The enhanced drug deposition in tumors that received laser treatment could be due to a combination of enhanced light-triggered drug release, phototherapy-induced dilation of blood vessels, and hyperthermia-mediated vascular permeabilization. The improved tumoral drug uptake is assumed to overcome the drug loss that occurs due to drug efflux attributed to P-gp and helps in overcoming Dox-resistance as observed in the anti-cancer efficacy study **(Fig. 4)**.

**Figure 5:**
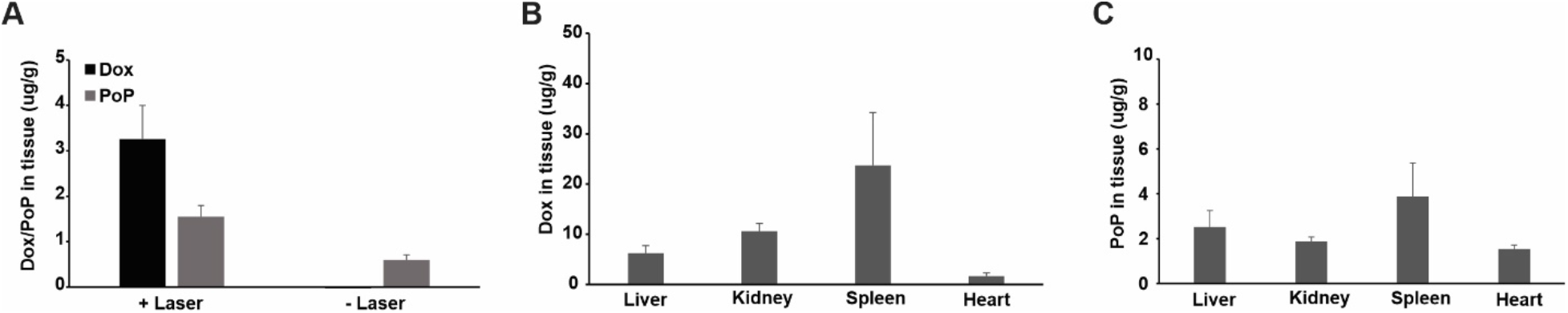
Tumor growth inhibition of Dox-resistant A2780 cells. A dual tumor model of nude mice bearing A2780-R tumors was treated with LC-Dox-PoP liposomes with 4 mg/kg Dox dose. Tumors that received laser treatment were treated using a 665 nm laser with a laser fluence rate of 200 mW/cm^2^ with a total fluence of 250 J/cm^2^. (A) Tumoral uptake of Dox/PoP with or without laser irradiation (B) Dox in tissue (C) PoP in tissue. Each data point shows the mean and ±SD of n=5 mice data obtained at each time point.

## 4. Conclusion

In this study, we demonstrated how phototherapy could be an effective tool to reverse MDR in cancer cells. We systematically studied the effect of phototherapy on A2780 and A2780-R cells using LC-Dox-PoP liposomes in vitro and in vivo. An in vitro study on both types of cells using free Dox and LC-Dox-PoP liposomes without phototherapy showed a significant difference in cell viability measured using the XTT assay. It was observed that A2780 cells showed around 20% cell viability, whereas A2780-R cells showed cell viability of above 60% and 70%, respectively, with free Dox and LC-Dox-PoP without phototherapy. Interestingly, we observed almost equivalent % cell viability of both cell types when treated with the same dose of LC-Dox-PoP liposomes with phototherapy. To examine this result further, we conducted another in vitro study on both these cell lines using empty PoP liposomes, LC-Dox-PoP liposomes, and free Dox with phototherapy. We found that LC-Dox-PoP liposomes with phototherapy using a total fluence of 10 J/cm^2^ showed similar %cell viability of both cell types. To investigate the impact of phototherapy on Dox-resistant tumors, an in vivo study was performed on nude mice bearing A2780-R cancer xenografts. This study illustrated faster regression of Dox-resistant tumors with phototherapy when administered with 4 mg/kg LC-Dox-PoP followed by phototherapy using a 665 nm laser with 200 mW/cm^2^ with a total fluence of 250 J/cm^2^ at 1 hour DLI. This aids in validating our hypothesis that phototherapy could pose to be a useful measure in curbing drug resistance in tumors. To corroborate the results obtained from the survival study, the biodistribution of Dox in tumors with and without phototherapy was assessed. A dual-tumor nude mouse model with A2780-R cells when administered with LC-Dox-PoP liposomes followed by phototherapy in one tumor demonstrated appreciably enhanced accumulation of Dox and PoP in tumors that received phototreatment. Synchronically, this data assists in understanding how phototherapy leads to improved tumoral drug uptake and faster tumor shrinkage. In the future, it would be interesting to study the impact of phototherapy on other types of drug resistance in other tumor models.

## 5. Acknowledgments

This study was supported by the National Institutes of Health (R01EB017270 and R01CA243164). J.F.L. holds interest in POP Biotechnologies.

